## supplementary information for "Do Larger Models Really Win in Drug Discovery? A Benchmark Assessment of Model Scaling in AI Driven Molecular Property and Activity Prediction"

### Supplementary Note 1: Evidence Index

This supplementary file is intentionally concise. The full benchmark evidence is provided as machine readable CSV files rather than as oversized typeset tables. The principal evidence files are:

- Integrated family best table for 78 split and task entries.
- Endpoint class definitions used in the manuscript.
- Family level winner counts by endpoint class.
- Winner counts by split protocol, endpoint class and family.
- Prevalence aware hERG/KCNH2, PXR/NR1I2, DRD2 and EGFR leader table with lift over prevalence.
- Summary and complete paired bootstrap test tables.
- Top K enrichment summaries.
- Calibration summaries.
- Plotting ready data regime scaling table.
- LLM-SAR with versus without learned SAR knowledge summary.
- Checksum manifests for raw prediction evidence.

### Supplementary Note 2: Split Sensitivity Matrix

The structure separated split is constructed from molecular fingerprints rather than from labels. For molecule $i$, let $x_i \in \{0,1\}^{2048}$ denote its ECFP4 fingerprint. TruncatedSVD maps $x_i$ to a lower dimensional vector $z_i$. MiniBatchKMeans then assigns each molecule to a chemical space cluster by minimizing

$$\sum_i \|z_i - \mu_{c_i}\|_2^2,$$

where $c_i$ is the cluster assignment and $\mu_{c_i}$ is the corresponding cluster centroid. Whole clusters, rather than individual molecules, are assigned to one of five folds. This keeps local chemical neighbourhoods within the same fold and reduces near neighbour leakage between training and held out molecules.

The split sensitivity analysis compares structure separated, random and Murcko scaffold five fold CV. The main text reports family winner counts by split and endpoint class, together with the split level family winner overview. The supplementary material therefore focuses on complete endpoint matrices at the individual model level for random and Murcko scaffold CV.

### Individual Model by Endpoint Matrices for Random and Murcko Splits

The main text reports the individual model by endpoint matrices under structure separated CV. For completeness, the corresponding random and Murcko scaffold split matrices are reported below using the same endpoint classes and highlighting rule. Classification tables report PR-AUC / ROC-AUC within each endpoint entry, and ADME related regression tables report MAE / Pearson. The larger toxicity related classification panel is split into general toxicity and safety liability endpoints, Tox21 nuclear receptor assays and Tox21 stress response assays. Rankings are computed independently for each metric within each endpoint: the first ranked value is bold and underlined, and the second and third ranked values are underlined.

#### Random five fold CV

Table 1: Random 5-fold CV model-by-endpoint matrix for ADME-related classification endpoints. Each endpoint entry reports PR-AUC / ROC-AUC. Rankings are computed independently for each metric within each endpoint: the first ranked value is bold and underlined, and the second and third ranked values are underlined.

| Family | Model | BBB | CYP3A4 | PXR/NR1I2 |
|---|---|---|---|---|
| ML | ExtraTrees(ECFP4) | 0.963 / 0.919 | 0.847 / 0.885 | 0.973 / 0.922 |
| ML | ExtraTrees(ECFP6) | 0.961 / 0.911 | 0.846 / 0.884 | 0.972 / 0.921 |
| ML | ExtraTrees(RDKit desc.) | 0.971 / **0.934** | 0.836 / 0.883 | 0.974 / 0.924 |
| ML | ExtraTrees(MACCS) | 0.963 / 0.919 | 0.782 / 0.848 | 0.966 / 0.906 |
| ML | RF(ECFP4) | 0.971 / 0.926 | 0.852 / 0.887 | 0.978 / 0.927 |
| ML | RF(ECFP6) | 0.966 / 0.915 | 0.849 / 0.884 | 0.976 / 0.924 |
| ML | RF(RDKit desc.) | 0.973 / 0.932 | 0.845 / 0.887 | 0.977 / 0.926 |
| ML | RF(MACCS) | 0.972 / 0.929 | 0.806 / 0.861 | 0.973 / 0.916 |
| ML | GBDT(ECFP4) | 0.962 / 0.901 | 0.839 / 0.875 | 0.966 / 0.909 |
| ML | GBDT(ECFP6) | 0.963 / 0.903 | 0.833 / 0.871 | 0.967 / 0.905 |
| ML | GBDT(RDKit desc.) | 0.969 / 0.922 | 0.848 / 0.891 | 0.973 / 0.919 |
| ML | GBDT(MACCS) | 0.966 / 0.913 | 0.800 / 0.855 | 0.971 / 0.909 |
| ML | LR(ECFP4) | 0.951 / 0.879 | 0.765 / 0.835 | 0.952 / 0.874 |
| ML | LR(ECFP6) | 0.947 / 0.870 | 0.770 / 0.832 | 0.946 / 0.865 |
| ML | LR(RDKit desc.) | 0.954 / 0.892 | 0.800 / 0.859 | 0.957 / 0.872 |
| ML | LR(MACCS) | 0.959 / 0.895 | 0.755 / 0.826 | 0.960 / 0.878 |
| GNN | Ligandformer | 0.962 / 0.910 | 0.847 / 0.890 | 0.968 / 0.907 |
| GNN | GAT | 0.967 / 0.918 | 0.854 / 0.893 | 0.971 / 0.907 |
| GNN | GCN | 0.964 / 0.910 | 0.845 / 0.889 | 0.963 / 0.889 |
| GNN | GIN | 0.963 / 0.911 | 0.859 / 0.899 | 0.973 / 0.914 |
| Sequence | ChemBERTa | 0.967 / 0.919 | 0.845 / 0.887 | 0.963 / 0.883 |
| Sequence | ChemBERTa2 | 0.971 / 0.923 | 0.868 / 0.905 | 0.975 / 0.916 |
| Sequence | MoLFormer | **0.974** / 0.927 | **0.873** / **0.908** | **0.981** / **0.937** |
| LLM-SAR | GPT5.5-SAR | 0.896 / 0.757 | 0.605 / 0.732 | 0.793 / 0.503 |
| LLM-SAR | GPT5.5-SAR + knowledge | 0.914 / 0.799 | 0.616 / 0.749 | 0.858 / 0.627 |
| LLM-SAR | Opus4.7-SAR | 0.895 / 0.793 | 0.612 / 0.733 | 0.781 / 0.519 |
| LLM-SAR | Opus4.7-SAR + knowledge | 0.933 / 0.834 | 0.674 / 0.783 | 0.853 / 0.653 |

Table 2: Random 5-fold CV model-by-endpoint matrix for ADME-related regression endpoints. Each endpoint entry reports MAE / Pearson. Lower MAE is better; higher Pearson is better. Rankings are computed independently for each metric within each endpoint: the first ranked value is bold and underlined, and the second and third ranked values are underlined.

| Family | Model | Caco2 | Lipophilicity | Solubility |
|---|---|---|---|---|
| ML | ExtraTrees(ECFP4) | 0.412 / 0.713 | 0.776 / 0.588 | 1.190 / 0.728 |
| ML | ExtraTrees(ECFP6) | 0.426 / 0.691 | 0.840 / 0.521 | 1.209 / 0.724 |
| ML | ExtraTrees(RDKit desc.) | <u>**0.293**</u> / <u>**0.869**</u> | <u>0.485</u> / <u>0.839</u> | <u>**0.653**</u> / <u>**0.909**</u> |
| ML | ExtraTrees(MACCS) | 0.363 / 0.782 | 0.771 / 0.601 | 1.100 / 0.765 |
| ML | RF(ECFP4) | 0.332 / 0.830 | 0.615 / 0.736 | 0.955 / 0.825 |
| ML | RF(ECFP6) | 0.340 / 0.825 | 0.646 / 0.710 | 0.961 / 0.824 |
| ML | RF(RDKit desc.) | <u>0.306</u> / <u>0.863</u> | 0.520 / 0.821 | <u>0.695</u> / <u>0.901</u> |
| ML | RF(MACCS) | 0.315 / 0.851 | 0.606 / 0.746 | 0.896 / 0.845 |
| ML | GBDT(ECFP4) | 0.341 / 0.822 | 0.669 / 0.713 | 1.120 / 0.783 |
| ML | GBDT(ECFP6) | 0.354 / 0.805 | 0.688 / 0.686 | 1.118 / 0.783 |
| ML | GBDT(RDKit desc.) | <u>0.298</u> / <u>0.868</u> | 0.520 / 0.819 | 0.726 / 0.897 |
| ML | GBDT(MACCS) | 0.324 / 0.842 | 0.656 / 0.712 | 0.991 / 0.823 |
| ML | Ridge(ECFP4) | 0.427 / 0.722 | 0.957 / 0.549 | 1.221 / 0.745 |
| ML | Ridge(ECFP6) | 0.422 / 0.730 | 0.925 / 0.569 | 1.234 / 0.744 |
| ML | Ridge(RDKit desc.) | 0.345 / 0.826 | 0.621 / 0.736 | 0.947 / 0.828 |
| ML | Ridge(MACCS) | 0.406 / 0.755 | 0.756 / 0.609 | 1.154 / 0.767 |
| Sequence | ChemBERTa | 0.625 / 0.295 | 0.640 / 0.765 | 0.814 / 0.882 |
| Sequence | ChemBERTa2 | 0.494 / 0.654 | <u>0.507</u> / <u>0.839</u> | 0.752 / 0.893 |
| Sequence | MoLFormer | 0.333 / 0.846 | <u>**0.436**</u> / <u>**0.870**</u> | <u>0.698</u> / <u>0.901</u> |
| LLM-SAR | GPT5.5-SAR | 0.646 / 0.154 | 1.371 / 0.405 | 2.287 / 0.564 |
| LLM-SAR | GPT5.5-SAR + knowledge | 0.562 / 0.486 | 2.096 / 0.458 | 1.731 / 0.595 |
| LLM-SAR | Opus4.7-SAR | 0.890 / 0.547 | 1.468 / 0.422 | 1.826 / 0.669 |
| LLM-SAR | Opus4.7-SAR + knowledge | 1.075 / 0.646 | 1.437 / 0.559 | 2.120 / 0.726 |

Table 3: Random 5-fold CV model-by-endpoint matrix for general toxicity and safety liability endpoints. Each endpoint entry reports PR-AUC / ROC-AUC. Rankings are computed independently for each metric within each endpoint: the first ranked value is bold and underlined, and the second and third ranked values are underlined.

| Family | Model | AMES | DILI | TDC hERG | hERG/KCNH2 | DRD2 |
|---|---|---|---|---|---|---|
| ML | ExtraTrees(ECFP4) | 0.921 / 0.908 | 0.890 / 0.893 | 0.907 / 0.846 | 0.885 / 0.949 | 0.991 / 0.953 |
| ML | ExtraTrees(ECFP6) | 0.920 / 0.906 | 0.881 / 0.886 | 0.908 / 0.846 | 0.883 / 0.948 | 0.991 / 0.953 |
| ML | ExtraTrees(RDKit desc.) | **0.921** / 0.909 | 0.913 / 0.909 | 0.925 / 0.875 | **0.892** / **0.953** | 0.988 / 0.938 |
| ML | ExtraTrees(MACCS) | 0.884 / 0.885 | 0.887 / 0.891 | 0.909 / 0.853 | 0.839 / 0.930 | 0.981 / 0.911 |
| ML | RF(ECFP4) | 0.917 / 0.905 | 0.885 / 0.887 | 0.907 / 0.847 | 0.886 / 0.949 | 0.993 / **0.958** |
| ML | RF(ECFP6) | 0.914 / 0.902 | 0.882 / 0.883 | 0.905 / 0.845 | 0.884 / 0.949 | **0.993** / 0.957 |
| ML | RF(RDKit desc.) | 0.919 / **0.911** | **0.915** / 0.908 | 0.933 / **0.882** | 0.882 / 0.949 | 0.991 / 0.942 |
| ML | RF(MACCS) | 0.908 / 0.898 | 0.893 / 0.894 | 0.918 / 0.860 | 0.861 / 0.939 | 0.989 / 0.934 |
| ML | GBDT(ECFP4) | 0.886 / 0.872 | 0.842 / 0.848 | 0.881 / 0.815 | 0.824 / 0.919 | 0.990 / 0.937 |
| ML | GBDT(ECFP6) | 0.884 / 0.867 | 0.841 / 0.848 | 0.892 / 0.831 | 0.827 / 0.918 | 0.991 / 0.939 |
| ML | GBDT(RDKit desc.) | 0.902 / 0.889 | 0.880 / 0.889 | 0.905 / 0.845 | 0.841 / 0.929 | 0.989 / 0.929 |
| ML | GBDT(MACCS) | 0.890 / 0.877 | 0.845 / 0.866 | 0.900 / 0.834 | 0.766 / 0.895 | 0.985 / 0.904 |
| ML | LR(ECFP4) | 0.789 / 0.784 | 0.838 / 0.842 | 0.868 / 0.796 | 0.726 / 0.884 | 0.982 / 0.900 |
| ML | LR(ECFP6) | 0.806 / 0.791 | 0.810 / 0.828 | 0.867 / 0.786 | 0.716 / 0.879 | 0.983 / 0.899 |
| ML | LR(RDKit desc.) | 0.853 / 0.842 | 0.821 / 0.822 | 0.869 / 0.799 | 0.621 / 0.850 | 0.977 / 0.859 |
| ML | LR(MACCS) | 0.831 / 0.823 | 0.808 / 0.809 | 0.831 / 0.749 | 0.596 / 0.818 | 0.975 / 0.846 |
| GNN | Ligandformer | 0.906 / 0.892 | 0.855 / 0.876 | 0.907 / 0.843 | 0.866 / 0.936 | 0.988 / 0.927 |
| GNN | GAT | 0.905 / 0.891 | 0.868 / 0.893 | 0.879 / 0.815 | 0.866 / 0.937 | 0.990 / 0.934 |
| GNN | GCN | 0.902 / 0.890 | 0.832 / 0.852 | 0.893 / 0.821 | 0.860 / 0.937 | 0.989 / 0.931 |
| GNN | GIN | 0.910 / 0.899 | 0.818 / 0.847 | 0.902 / 0.834 | 0.889 / 0.946 | 0.989 / 0.936 |
| Sequence | ChemBERTa | 0.894 / 0.879 | 0.834 / 0.843 | 0.899 / 0.831 | 0.825 / 0.923 | 0.986 / 0.912 |
| Sequence | ChemBERTa2 | 0.903 / 0.888 | 0.883 / 0.883 | 0.930 / 0.876 | 0.863 / 0.937 | 0.990 / 0.937 |
| Sequence | MoLFormer | 0.842 / 0.830 | 0.905 / **0.913** | 0.933 / 0.878 | 0.877 / 0.942 | 0.992 / 0.947 |
| LLM-SAR | GPT5.5-SAR | 0.642 / 0.614 | 0.691 / 0.681 | 0.934 / 0.809 | 0.286 / 0.618 | 0.920 / 0.612 |
| LLM-SAR | GPT5.5-SAR + knowledge | 0.703 / 0.668 | 0.750 / 0.758 | **0.943** / 0.825 | 0.356 / 0.694 | 0.925 / 0.646 |
| LLM-SAR | Opus4.7-SAR | 0.751 / 0.726 | 0.554 / 0.565 | 0.913 / 0.844 | 0.368 / 0.703 | 0.919 / 0.604 |
| LLM-SAR | Opus4.7-SAR + knowledge | 0.792 / 0.763 | 0.778 / 0.780 | 0.903 / 0.838 | 0.445 / 0.736 | 0.955 / 0.753 |

Table 4: Random 5-fold CV model-by-endpoint matrix for Tox21 nuclear receptor assays. Each endpoint entry reports PR-AUC / ROC-AUC. Rankings are computed independently for each metric within each endpoint: the first ranked value is bold and underlined, and the second and third ranked values are underlined.

| Family | Model | NR-AR | NR-AR-LBD | NR-AhR | NR-Aromatase | NR-ER | NR-ER-LBD | NR-PPAR-$\gamma$ |
|---|---|---|---|---|---|---|---|---|
| ML | ExtraTrees(ECFP4) | 0.474 / 0.785 | 0.648 / 0.867 | 0.607 / 0.892 | 0.384 / 0.815 | 0.362 / 0.708 | 0.400 / 0.804 | 0.294 / 0.837 |
| ML | ExtraTrees(ECFP6) | 0.483 / 0.782 | 0.648 / 0.863 | 0.607 / 0.894 | 0.419 / 0.826 | 0.363 / 0.704 | 0.396 / 0.800 | 0.279 / 0.831 |
| ML | ExtraTrees(RDKit desc.) | 0.512 / 0.796 | **0.654** / **0.894** | 0.640 / 0.903 | 0.443 / 0.859 | 0.406 / 0.720 | 0.476 / 0.829 | **0.327** / 0.856 |
| ML | ExtraTrees(MACCS) | 0.434 / 0.773 | 0.565 / 0.858 | 0.548 / 0.873 | 0.329 / 0.826 | 0.323 / 0.693 | 0.346 / 0.787 | 0.209 / 0.780 |
| ML | RF(ECFP4) | 0.523 / 0.789 | 0.651 / 0.869 | 0.622 / 0.896 | 0.403 / 0.822 | 0.419 / 0.722 | 0.457 / 0.817 | 0.293 / 0.842 |
| ML | RF(ECFP6) | 0.528 / 0.781 | 0.646 / 0.861 | 0.615 / 0.896 | 0.434 / 0.830 | 0.409 / 0.717 | 0.447 / 0.815 | 0.277 / 0.834 |
| ML | RF(RDKit desc.) | **0.537** / 0.812 | 0.644 / 0.880 | **0.645** / **0.905** | **0.444** / **0.860** | **0.469** / 0.733 | **0.521** / **0.839** | 0.314 / 0.854 |
| ML | RF(MACCS) | 0.506 / 0.784 | 0.598 / 0.863 | 0.595 / 0.892 | 0.377 / 0.838 | 0.397 / 0.712 | 0.432 / 0.816 | 0.214 / 0.807 |
| ML | GBDT(ECFP4) | 0.507 / 0.773 | 0.598 / 0.836 | 0.568 / 0.879 | 0.371 / 0.812 | 0.412 / 0.694 | 0.467 / 0.778 | 0.241 / 0.774 |
| ML | GBDT(ECFP6) | 0.502 / 0.757 | 0.606 / 0.847 | 0.546 / 0.869 | 0.376 / 0.794 | 0.410 / 0.696 | 0.454 / 0.769 | 0.238 / 0.742 |
| ML | GBDT(RDKit desc.) | 0.523 / 0.785 | 0.602 / 0.876 | 0.634 / 0.901 | 0.417 / 0.844 | 0.458 / 0.713 | 0.506 / 0.819 | 0.314 / 0.849 |
| ML | GBDT(MACCS) | 0.511 / 0.768 | 0.579 / 0.823 | 0.585 / 0.887 | 0.334 / 0.824 | 0.400 / 0.707 | 0.419 / 0.772 | 0.253 / 0.798 |
| ML | LR(ECFP4) | 0.386 / 0.718 | 0.536 / 0.832 | 0.413 / 0.799 | 0.279 / 0.758 | 0.252 / 0.627 | 0.330 / 0.757 | 0.199 / 0.746 |
| ML | LR(ECFP6) | 0.411 / 0.704 | 0.568 / 0.822 | 0.395 / 0.783 | 0.277 / 0.731 | 0.225 / 0.603 | 0.305 / 0.719 | 0.218 / 0.728 |
| ML | LR(RDKit desc.) | 0.434 / 0.762 | 0.415 / 0.835 | 0.575 / 0.888 | 0.217 / 0.798 | 0.339 / 0.713 | 0.281 / 0.796 | 0.163 / 0.785 |
| ML | LR(MACCS) | 0.431 / 0.786 | 0.386 / 0.827 | 0.504 / 0.867 | 0.219 / 0.806 | 0.328 / 0.686 | 0.232 / 0.749 | 0.146 / 0.782 |
| GNN | Ligandformer | 0.501 / 0.806 | 0.548 / 0.874 | 0.555 / 0.883 | 0.339 / 0.848 | 0.436 / 0.723 | 0.464 / 0.827 | 0.303 / 0.852 |
| GNN | GAT | 0.453 / 0.793 | 0.479 / 0.878 | 0.586 / 0.891 | 0.338 / 0.828 | 0.384 / 0.718 | 0.389 / 0.810 | 0.235 / 0.845 |
| GNN | GCN | 0.467 / 0.797 | 0.582 / 0.880 | 0.580 / 0.887 | 0.352 / 0.829 | 0.396 / 0.718 | 0.381 / 0.820 | 0.257 / 0.834 |
| GNN | GIN | 0.475 / 0.804 | 0.566 / 0.879 | 0.580 / 0.894 | 0.335 / 0.845 | 0.427 / 0.717 | 0.457 / 0.826 | 0.307 / **0.857** |
| Sequence | ChemBERTa | 0.518 / 0.815 | 0.567 / 0.861 | 0.568 / 0.887 | 0.316 / 0.834 | 0.417 / 0.731 | 0.447 / 0.818 | 0.259 / 0.786 |
| Sequence | ChemBERTa2 | 0.534 / **0.821** | 0.592 / 0.873 | 0.625 / 0.902 | 0.355 / 0.855 | 0.454 / **0.740** | 0.477 / 0.833 | 0.291 / 0.838 |
| Sequence | MoLFormer | 0.401 / 0.812 | 0.442 / 0.838 | 0.642 / 0.901 | 0.284 / 0.822 | 0.387 / 0.709 | 0.446 / 0.813 | 0.217 / 0.782 |
| LLM-SAR | GPT5.5-SAR | 0.049 / 0.513 | 0.038 / 0.501 | 0.229 / 0.732 | 0.175 / 0.762 | 0.176 / 0.618 | 0.070 / 0.608 | 0.090 / 0.686 |
| LLM-SAR | GPT5.5-SAR + knowledge | 0.073 / 0.696 | 0.064 / 0.712 | 0.250 / 0.764 | 0.187 / 0.783 | 0.189 / 0.653 | 0.080 / 0.661 | 0.093 / 0.719 |
| LLM-SAR | Opus4.7-SAR | 0.056 / 0.618 | 0.058 / 0.614 | 0.231 / 0.726 | 0.198 / 0.753 | 0.194 / 0.626 | 0.108 / 0.671 | 0.178 / 0.720 |
| LLM-SAR | Opus4.7-SAR + knowledge | 0.156 / 0.750 | 0.196 / 0.786 | 0.358 / 0.789 | 0.215 / 0.779 | 0.211 / 0.645 | 0.101 / 0.670 | 0.140 / 0.714 |

Table 5: Random 5-fold CV model-by-endpoint matrix for Tox21 stress response assays. Each endpoint entry reports PR-AUC / ROC-AUC. Rankings are computed independently for each metric within each endpoint: the first ranked value is bold and underlined, and the second and third ranked values are underlined.

| Family | Model | SR-ARE | SR-ATAD5 | SR-HSE | SR-MMP | SR-p53 |
|---|---|---|---|---|---|---|
| ML | ExtraTrees(ECFP4) | 0.500 / 0.804 | 0.315 / 0.843 | 0.272 / 0.774 | 0.655 / 0.879 | 0.413 / 0.855 |
| ML | ExtraTrees(ECFP6) | 0.502 / 0.801 | 0.321 / 0.844 | 0.279 / 0.780 | 0.655 / 0.879 | 0.397 / 0.847 |
| ML | ExtraTrees(RDKit desc.) | 0.552 / 0.839 | **0.350** / **0.877** | **0.388** / 0.809 | 0.762 / 0.928 | **0.484** / **0.886** |
| ML | ExtraTrees(MACCS) | 0.493 / 0.809 | 0.272 / 0.841 | 0.238 / 0.755 | 0.667 / 0.887 | 0.357 / 0.838 |
| ML | RF(ECFP4) | 0.515 / 0.810 | 0.334 / 0.861 | 0.289 / 0.778 | 0.670 / 0.881 | 0.414 / 0.852 |
| ML | RF(ECFP6) | 0.516 / 0.807 | 0.332 / 0.855 | 0.296 / 0.781 | 0.664 / 0.881 | 0.400 / 0.841 |
| ML | RF(RDKit desc.) | **0.554** / **0.841** | 0.338 / 0.875 | 0.379 / 0.804 | **0.770** / **0.929** | 0.450 / 0.882 |
| ML | RF(MACCS) | 0.517 / 0.822 | 0.306 / 0.875 | 0.285 / 0.773 | 0.692 / 0.900 | 0.380 / 0.856 |
| ML | GBDT(ECFP4) | 0.456 / 0.763 | 0.274 / 0.817 | 0.273 / 0.737 | 0.629 / 0.859 | 0.328 / 0.802 |
| ML | GBDT(ECFP6) | 0.442 / 0.749 | 0.271 / 0.798 | 0.252 / 0.713 | 0.602 / 0.851 | 0.335 / 0.791 |
| ML | GBDT(RDKit desc.) | 0.544 / 0.833 | 0.347 / 0.850 | 0.364 / 0.793 | 0.753 / 0.923 | 0.398 / 0.863 |
| ML | GBDT(MACCS) | 0.463 / 0.789 | 0.287 / 0.811 | 0.312 / 0.755 | 0.646 / 0.886 | 0.316 / 0.824 |
| ML | LR(ECFP4) | 0.334 / 0.681 | 0.255 / 0.751 | 0.169 / 0.655 | 0.479 / 0.792 | 0.236 / 0.715 |
| ML | LR(ECFP6) | 0.297 / 0.651 | 0.249 / 0.766 | 0.154 / 0.652 | 0.465 / 0.779 | 0.234 / 0.725 |
| ML | LR(RDKit desc.) | 0.416 / 0.783 | 0.191 / 0.807 | 0.271 / 0.778 | 0.609 / 0.885 | 0.268 / 0.828 |
| ML | LR(MACCS) | 0.371 / 0.752 | 0.167 / 0.780 | 0.239 / 0.729 | 0.534 / 0.853 | 0.232 / 0.794 |
| GNN | Ligandformer | 0.510 / 0.813 | 0.256 / 0.851 | 0.277 / 0.789 | 0.702 / 0.916 | 0.389 / 0.854 |
| GNN | GAT | 0.503 / 0.807 | 0.233 / 0.842 | 0.328 / 0.785 | 0.689 / 0.901 | 0.360 / 0.847 |
| GNN | GCN | 0.492 / 0.802 | 0.243 / 0.843 | 0.304 / 0.792 | 0.718 / 0.903 | 0.379 / 0.855 |
| GNN | GIN | 0.538 / 0.821 | 0.293 / 0.862 | 0.349 / **0.813** | 0.741 / 0.914 | 0.392 / 0.870 |
| Sequence | ChemBERTa | 0.481 / 0.805 | 0.306 / 0.822 | 0.267 / 0.772 | 0.684 / 0.906 | 0.324 / 0.820 |
| Sequence | ChemBERTa2 | 0.529 / 0.822 | 0.309 / 0.847 | 0.319 / 0.796 | 0.748 / 0.921 | 0.398 / 0.860 |
| Sequence | MoLFormer | 0.518 / 0.826 | 0.310 / 0.843 | 0.317 / 0.803 | 0.741 / 0.917 | 0.234 / 0.760 |
| LLM-SAR | GPT5.5-SAR | 0.232 / 0.622 | 0.060 / 0.625 | 0.084 / 0.573 | 0.242 / 0.632 | 0.099 / 0.641 |
| LLM-SAR | GPT5.5-SAR + knowledge | 0.271 / 0.686 | 0.069 / 0.646 | 0.104 / 0.611 | 0.335 / 0.760 | 0.126 / 0.703 |
| LLM-SAR | Opus4.7-SAR | 0.201 / 0.535 | 0.068 / 0.617 | 0.128 / 0.662 | 0.279 / 0.718 | 0.131 / 0.619 |
| LLM-SAR | Opus4.7-SAR + knowledge | 0.321 / 0.706 | 0.066 / 0.620 | 0.154 / 0.699 | 0.419 / 0.797 | 0.178 / 0.715 |

Table 6: Random 5-fold CV model-by-endpoint matrix for bioactivity-related classification endpoints. Each endpoint entry reports PR-AUC / ROC-AUC. Rankings are computed independently for each metric within each endpoint: the first ranked value is bold and underlined, and the second and third ranked values are underlined.

| Family | Model | EGFR | anti-TB H37Rv | antimalaria Pf. 3D7/Dd2 |
|---|---|---|---|---|
| ML | ExtraTrees(ECFP4) | 0.989 / 0.969 | 0.584 / 0.858 | 0.968 / **0.965** |
| ML | ExtraTrees(ECFP6) | 0.990 / 0.970 | 0.597 / 0.859 | 0.966 / 0.964 |
| ML | ExtraTrees(RDKit desc.) | 0.986 / 0.958 | 0.579 / 0.856 | 0.968 / 0.965 |
| ML | ExtraTrees(MACCS) | 0.979 / 0.946 | 0.474 / 0.804 | 0.957 / 0.956 |
| ML | RF(ECFP4) | 0.992 / 0.972 | **0.609** / **0.868** | 0.966 / 0.963 |
| ML | RF(ECFP6) | **0.992** / **0.972** | 0.607 / 0.866 | 0.964 / 0.962 |
| ML | RF(RDKit desc.) | 0.988 / 0.959 | 0.592 / 0.863 | 0.963 / 0.959 |
| ML | RF(MACCS) | 0.986 / 0.957 | 0.529 / 0.841 | 0.958 / 0.955 |
| ML | GBDT(ECFP4) | 0.984 / 0.948 | 0.531 / 0.833 | 0.953 / 0.948 |
| ML | GBDT(ECFP6) | 0.985 / 0.952 | 0.535 / 0.836 | 0.949 / 0.944 |
| ML | GBDT(RDKit desc.) | 0.984 / 0.945 | 0.532 / 0.836 | 0.956 / 0.954 |
| ML | GBDT(MACCS) | 0.978 / 0.928 | 0.458 / 0.812 | 0.939 / 0.935 |
| ML | LR(ECFP4) | 0.970 / 0.912 | 0.346 / 0.771 | 0.924 / 0.929 |
| ML | LR(ECFP6) | 0.971 / 0.915 | 0.386 / 0.787 | 0.927 / 0.930 |
| ML | LR(RDKit desc.) | 0.964 / 0.887 | 0.362 / 0.760 | 0.920 / 0.922 |
| ML | LR(MACCS) | 0.963 / 0.883 | 0.320 / 0.749 | 0.866 / 0.871 |
| GNN | Ligandformer | 0.986 / 0.953 | 0.519 / 0.828 | 0.960 / 0.955 |
| GNN | GAT | 0.987 / 0.956 | 0.536 / 0.845 | 0.953 / 0.951 |
| GNN | GCN | 0.986 / 0.954 | 0.533 / 0.842 | 0.949 / 0.949 |
| GNN | GIN | 0.988 / 0.960 | 0.567 / 0.850 | 0.961 / 0.959 |
| Sequence | ChemBERTa | 0.982 / 0.941 | 0.530 / 0.827 | 0.950 / 0.943 |
| Sequence | ChemBERTa2 | 0.987 / 0.954 | 0.546 / 0.843 | 0.958 / 0.952 |
| Sequence | MoLFormer | 0.989 / 0.961 | 0.534 / 0.840 | **0.968** / 0.963 |
| LLM-SAR | GPT5.5-SAR | 0.858 / 0.675 | 0.126 / 0.473 | 0.611 / 0.630 |
| LLM-SAR | GPT5.5-SAR + knowledge | 0.865 / 0.692 | 0.163 / 0.578 | 0.672 / 0.699 |
| LLM-SAR | Opus4.7-SAR | 0.860 / 0.670 | 0.146 / 0.451 | 0.712 / 0.681 |
| LLM-SAR | Opus4.7-SAR + knowledge | 0.923 / 0.776 | 0.234 / 0.631 | 0.784 / 0.801 |

Table 7: Murcko scaffold 5-fold CV model-by-endpoint matrix for ADME-related classification endpoints. Each endpoint entry reports PR-AUC / ROC-AUC. Rankings are computed independently for each metric within each endpoint: the first ranked value is bold and underlined, and the second and third ranked values are underlined.

| Family | Model | BBB | CYP3A4 | PXR/NR1I2 |
|---|---|---|---|---|
| ML | ExtraTrees(ECFP4) | 0.952 / 0.887 | 0.747 / 0.825 | 0.954 / 0.887 |
| ML | ExtraTrees(ECFP6) | 0.953 / 0.887 | 0.750 / 0.825 | 0.950 / 0.882 |
| ML | ExtraTrees(RDKit desc.) | **<u>0.965</u>** / <u>0.905</u> | 0.717 / 0.829 | <u>0.956</u> / <u>0.899</u> |
| ML | ExtraTrees(MACCS) | 0.959 / 0.887 | 0.668 / 0.774 | 0.946 / 0.868 |
| ML | RF(ECFP4) | 0.954 / 0.890 | 0.754 / 0.826 | 0.954 / 0.885 |
| ML | RF(ECFP6) | 0.952 / 0.883 | 0.757 / 0.827 | 0.949 / 0.883 |
| ML | RF(RDKit desc.) | <u>0.965</u> / <u>0.899</u> | 0.731 / 0.840 | <u>0.957</u> / <u>0.897</u> |
| ML | RF(MACCS) | 0.962 / 0.890 | 0.687 / 0.787 | 0.951 / 0.877 |
| ML | GBDT(ECFP4) | 0.941 / 0.860 | 0.741 / 0.823 | 0.940 / 0.869 |
| ML | GBDT(ECFP6) | 0.935 / 0.842 | 0.732 / 0.814 | 0.936 / 0.860 |
| ML | GBDT(RDKit desc.) | 0.950 / 0.879 | 0.747 / 0.851 | 0.949 / 0.888 |
| ML | GBDT(MACCS) | 0.944 / 0.867 | 0.670 / 0.788 | 0.935 / 0.847 |
| ML | LR(ECFP4) | 0.919 / 0.826 | 0.663 / 0.780 | 0.922 / 0.821 |
| ML | LR(ECFP6) | 0.912 / 0.804 | 0.673 / 0.781 | 0.919 / 0.825 |
| ML | LR(RDKit desc.) | 0.936 / 0.854 | 0.700 / 0.821 | 0.937 / 0.849 |
| ML | LR(MACCS) | 0.926 / 0.832 | 0.616 / 0.750 | 0.930 / 0.821 |
| GNN | Ligandformer | 0.955 / 0.880 | <u>0.782</u> / 0.862 | 0.950 / 0.883 |
| GNN | GAT | 0.953 / 0.877 | **<u>0.800</u>** / **<u>0.868</u>** | 0.956 / 0.893 |
| GNN | GCN | 0.941 / 0.868 | <u>0.790</u> / <u>0.865</u> | 0.938 / 0.848 |
| GNN | GIN | 0.949 / 0.876 | 0.771 / <u>0.863</u> | 0.953 / 0.875 |
| Sequence | ChemBERTa | 0.951 / 0.882 | 0.722 / 0.831 | 0.930 / 0.820 |
| Sequence | ChemBERTa2 | 0.953 / 0.887 | 0.763 / 0.856 | 0.955 / 0.884 |
| Sequence | MoLFormer | <u>0.964</u> / **<u>0.906</u>** | 0.729 / 0.806 | **<u>0.970</u>** / **<u>0.917</u>** |
| LLM-SAR | GPT5.5-SAR | 0.897 / 0.753 | 0.604 / 0.729 | 0.797 / 0.443 |
| LLM-SAR | GPT5.5-SAR + knowledge | 0.913 / 0.793 | 0.614 / 0.746 | 0.847 / 0.533 |
| LLM-SAR | Opus4.7-SAR | 0.895 / 0.784 | 0.617 / 0.739 | 0.782 / 0.435 |
| LLM-SAR | Opus4.7-SAR + knowledge | 0.929 / 0.822 | 0.650 / 0.778 | 0.825 / 0.515 |

**Murcko scaffold five fold CV**

Table 8: Murcko scaffold 5-fold CV model-by-endpoint matrix for ADME-related regression endpoints. Each endpoint entry reports MAE / Pearson. Lower MAE is better; higher Pearson is better. Rankings are computed independently for each metric within each endpoint: the first ranked value is bold and underlined, and the second and third ranked values are underlined.

| Family | Model | Caco2 | Lipophilicity | Solubility |
|---|---|---|---|---|
| ML | ExtraTrees(ECFP4) | 0.532 / 0.570 | 0.904 / 0.487 | 1.559 / 0.621 |
| ML | ExtraTrees(ECFP6) | 0.540 / 0.526 | 0.947 / 0.433 | 1.580 / 0.599 |
| ML | ExtraTrees(RDKit desc.) | 0.356 / 0.810 | 0.554 / 0.802 | **0.730** / **0.889** |
| ML | ExtraTrees(MACCS) | 0.486 / 0.644 | 0.941 / 0.439 | 1.295 / 0.674 |
| ML | RF(ECFP4) | 0.428 / 0.726 | 0.702 / 0.656 | 1.291 / 0.739 |
| ML | RF(ECFP6) | 0.435 / 0.726 | 0.734 / 0.623 | 1.280 / 0.738 |
| ML | RF(RDKit desc.) | 0.367 / 0.797 | 0.577 / 0.785 | 0.762 / 0.881 |
| ML | RF(MACCS) | 0.387 / 0.768 | 0.701 / 0.659 | 1.020 / 0.792 |
| ML | GBDT(ECFP4) | 0.404 / 0.753 | 0.719 / 0.657 | 1.296 / 0.731 |
| ML | GBDT(ECFP6) | 0.477 / 0.651 | 0.746 / 0.617 | 1.306 / 0.723 |
| ML | GBDT(RDKit desc.) | **0.356** / **0.811** | 0.557 / 0.796 | 0.759 / 0.883 |
| ML | GBDT(MACCS) | 0.383 / 0.783 | 0.711 / 0.660 | 1.074 / 0.795 |
| ML | Ridge(ECFP4) | 0.542 / 0.566 | 1.163 / 0.460 | 1.441 / 0.672 |
| ML | Ridge(ECFP6) | 0.522 / 0.611 | 1.065 / 0.489 | 1.408 / 0.678 |
| ML | Ridge(RDKit desc.) | 0.389 / 0.779 | 0.645 / 0.713 | 0.981 / 0.797 |
| ML | Ridge(MACCS) | 0.445 / 0.718 | 0.786 / 0.568 | 1.190 / 0.743 |
| Sequence | ChemBERTa | 0.627 / 0.258 | 0.658 / 0.745 | 0.889 / 0.856 |
| Sequence | ChemBERTa2 | 0.501 / 0.616 | 0.539 / 0.815 | 0.797 / 0.875 |
| Sequence | MoLFormer | 0.383 / 0.793 | **0.479** / **0.845** | 0.762 / 0.883 |
| LLM-SAR | GPT5.5-SAR | 0.646 / 0.151 | 1.371 / 0.400 | 2.287 / 0.537 |
| LLM-SAR | GPT5.5-SAR + knowledge | 0.605 / 0.399 | 2.059 / 0.443 | 1.763 / 0.573 |
| LLM-SAR | Opus4.7-SAR | 0.890 / 0.540 | 1.468 / 0.418 | 1.826 / 0.647 |
| LLM-SAR | Opus4.7-SAR + knowledge | 1.069 / 0.635 | 1.441 / 0.547 | 2.029 / 0.704 |

Table 9: Murcko scaffold 5-fold CV model-by-endpoint matrix for general toxicity and safety liability endpoints. Each endpoint entry reports PR-AUC / ROC-AUC. Rankings are computed independently for each metric within each endpoint: the first ranked value is bold and underlined, and the second and third ranked values are underlined.

| Family | Model | AMES | DILI | TDC hERG | hERG/KCNH2 | DRD2 |
|---|---|---|---|---|---|---|
| ML | ExtraTrees(ECFP4) | 0.843 / 0.825 | 0.808 / 0.836 | 0.894 / 0.809 | 0.793 / 0.909 | 0.986 / 0.922 |
| ML | ExtraTrees(ECFP6) | 0.839 / 0.821 | 0.805 / 0.827 | 0.897 / 0.814 | 0.785 / 0.905 | **0.987** / **0.927** |
| ML | ExtraTrees(RDKit desc.) | 0.844 / 0.827 | 0.853 / 0.861 | 0.913 / 0.868 | **0.812** / **0.919** | 0.979 / 0.891 |
| ML | ExtraTrees(MACCS) | 0.831 / 0.820 | 0.846 / 0.856 | 0.912 / 0.842 | 0.742 / 0.890 | 0.967 / 0.863 |
| ML | RF(ECFP4) | 0.849 / 0.826 | 0.808 / 0.835 | 0.889 / 0.805 | 0.787 / 0.906 | 0.986 / 0.923 |
| ML | RF(ECFP6) | 0.839 / 0.817 | 0.806 / 0.819 | 0.898 / 0.814 | 0.779 / 0.901 | 0.987 / 0.924 |
| ML | RF(RDKit desc.) | 0.839 / 0.822 | 0.847 / 0.852 | 0.912 / **0.868** | 0.788 / 0.907 | 0.980 / 0.888 |
| ML | RF(MACCS) | 0.843 / 0.829 | 0.849 / 0.857 | 0.912 / 0.844 | 0.762 / 0.894 | 0.974 / 0.876 |
| ML | GBDT(ECFP4) | 0.819 / 0.795 | 0.754 / 0.772 | 0.872 / 0.772 | 0.709 / 0.866 | 0.980 / 0.892 |
| ML | GBDT(ECFP6) | 0.820 / 0.792 | 0.724 / 0.768 | 0.875 / 0.781 | 0.708 / 0.869 | 0.981 / 0.892 |
| ML | GBDT(RDKit desc.) | 0.834 / 0.818 | 0.812 / 0.834 | 0.899 / 0.847 | 0.741 / 0.884 | 0.977 / 0.881 |
| ML | GBDT(MACCS) | 0.830 / 0.825 | 0.786 / 0.789 | 0.876 / 0.805 | 0.647 / 0.842 | 0.969 / 0.844 |
| ML | LR(ECFP4) | 0.695 / 0.681 | 0.797 / 0.804 | 0.873 / 0.767 | 0.604 / 0.829 | 0.960 / 0.824 |
| ML | LR(ECFP6) | 0.730 / 0.708 | 0.755 / 0.775 | 0.874 / 0.764 | 0.588 / 0.817 | 0.964 / 0.835 |
| ML | LR(RDKit desc.) | 0.794 / 0.778 | 0.811 / 0.820 | 0.848 / 0.782 | 0.541 / 0.800 | 0.962 / 0.816 |
| ML | LR(MACCS) | 0.772 / 0.765 | 0.723 / 0.690 | 0.853 / 0.774 | 0.508 / 0.771 | 0.959 / 0.802 |
| GNN | Ligandformer | 0.853 / 0.841 | 0.845 / 0.864 | 0.900 / 0.835 | 0.776 / 0.898 | 0.980 / 0.888 |
| GNN | GAT | 0.856 / 0.842 | 0.849 / 0.876 | 0.865 / 0.818 | 0.784 / 0.902 | 0.980 / 0.890 |
| GNN | GCN | 0.838 / 0.822 | 0.809 / 0.826 | 0.863 / 0.801 | 0.780 / 0.898 | 0.980 / 0.894 |
| GNN | GIN | **0.865** / **0.851** | 0.815 / 0.837 | 0.846 / 0.777 | 0.811 / 0.914 | 0.982 / 0.903 |
| Sequence | ChemBERTa | 0.829 / 0.810 | 0.786 / 0.807 | 0.879 / 0.812 | 0.721 / 0.874 | 0.973 / 0.859 |
| Sequence | ChemBERTa2 | 0.854 / 0.838 | 0.840 / 0.857 | 0.905 / 0.863 | 0.782 / 0.901 | 0.980 / 0.890 |
| Sequence | MoLFormer | 0.856 / 0.843 | **0.861** / **0.886** | 0.913 / 0.860 | 0.801 / 0.910 | 0.983 / 0.904 |
| LLM-SAR | GPT5.5-SAR | 0.654 / 0.624 | 0.704 / 0.657 | 0.928 / 0.810 | 0.296 / 0.637 | 0.915 / 0.634 |
| LLM-SAR | GPT5.5-SAR + knowledge | 0.701 / 0.650 | 0.736 / 0.722 | **0.938** / 0.825 | 0.344 / 0.691 | 0.921 / 0.673 |
| LLM-SAR | Opus4.7-SAR | 0.764 / 0.732 | 0.592 / 0.472 | 0.908 / 0.846 | 0.382 / 0.712 | 0.912 / 0.619 |
| LLM-SAR | Opus4.7-SAR + knowledge | 0.787 / 0.747 | 0.752 / 0.735 | 0.896 / 0.838 | 0.426 / 0.725 | 0.946 / 0.749 |

Table 10: Murcko scaffold 5-fold CV model-by-endpoint matrix for Tox21 nuclear receptor assays. Each endpoint entry reports PR-AUC / ROC-AUC. Rankings are computed independently for each metric within each endpoint: the first ranked value is bold and underlined, and the second and third ranked values are underlined.

| Family | Model | NR-AR | NR-AR-LBD | NR-AhR | NR-Aromatase | NR-ER | NR-ER-LBD | NR-PPAR-$\gamma$ |
|---|---|---|---|---|---|---|---|---|
| ML | ExtraTrees(ECFP4) | 0.475 / 0.788 | 0.566 / 0.862 | 0.494 / 0.870 | 0.211 / 0.736 | 0.245 / 0.644 | 0.262 / 0.740 | 0.248 / 0.803 |
| ML | ExtraTrees(ECFP6) | 0.493 / 0.785 | 0.558 / 0.856 | 0.511 / 0.871 | 0.219 / 0.758 | 0.255 / 0.649 | 0.274 / 0.763 | 0.246 / 0.803 |
| ML | ExtraTrees(RDKit desc.) | 0.518 / 0.782 | 0.564 / 0.864 | 0.490 / 0.857 | 0.260 / 0.792 | 0.372 / 0.702 | 0.412 / 0.815 | 0.288 / 0.847 |
| ML | ExtraTrees(MACCS) | 0.437 / 0.763 | 0.497 / 0.836 | 0.433 / 0.845 | 0.189 / 0.737 | 0.259 / 0.659 | 0.282 / 0.783 | 0.170 / 0.790 |
| ML | RF(ECFP4) | 0.543 / 0.782 | **0.580** / 0.862 | 0.526 / 0.874 | 0.227 / 0.741 | 0.276 / 0.651 | 0.300 / 0.753 | 0.264 / 0.821 |
| ML | RF(ECFP6) | 0.547 / 0.782 | 0.570 / 0.852 | 0.514 / 0.870 | 0.228 / 0.772 | 0.290 / 0.656 | 0.301 / 0.768 | 0.268 / 0.813 |
| ML | RF(RDKit desc.) | 0.545 / 0.788 | 0.571 / 0.870 | 0.482 / 0.862 | 0.283 / 0.788 | **0.426** / 0.706 | **0.442** / **0.835** | 0.289 / 0.833 |
| ML | RF(MACCS) | 0.513 / 0.767 | 0.549 / 0.853 | 0.484 / 0.864 | 0.204 / 0.739 | 0.312 / 0.678 | 0.337 / 0.792 | 0.188 / 0.789 |
| ML | GBDT(ECFP4) | 0.535 / 0.779 | 0.527 / 0.831 | 0.454 / 0.854 | 0.214 / 0.715 | 0.321 / 0.623 | 0.340 / 0.729 | 0.244 / 0.761 |
| ML | GBDT(ECFP6) | 0.538 / 0.764 | 0.542 / 0.822 | 0.419 / 0.831 | 0.206 / 0.733 | 0.290 / 0.627 | 0.344 / 0.745 | 0.195 / 0.722 |
| ML | GBDT(RDKit desc.) | 0.543 / 0.785 | 0.509 / 0.856 | 0.477 / 0.863 | 0.275 / 0.802 | 0.377 / 0.690 | 0.381 / 0.785 | 0.281 / 0.815 |
| ML | GBDT(MACCS) | 0.519 / 0.779 | 0.518 / 0.795 | 0.487 / 0.861 | 0.224 / 0.738 | 0.312 / 0.678 | 0.287 / 0.768 | 0.219 / 0.750 |
| ML | LR(ECFP4) | 0.413 / 0.748 | 0.483 / 0.826 | 0.301 / 0.745 | 0.143 / 0.657 | 0.196 / 0.574 | 0.208 / 0.687 | 0.190 / 0.740 |
| ML | LR(ECFP6) | 0.449 / 0.734 | 0.516 / 0.809 | 0.301 / 0.739 | 0.128 / 0.625 | 0.166 / 0.542 | 0.181 / 0.675 | 0.163 / 0.726 |
| ML | LR(RDKit desc.) | 0.363 / 0.752 | 0.381 / 0.803 | 0.473 / 0.856 | 0.188 / 0.747 | 0.259 / 0.665 | 0.199 / 0.766 | 0.169 / 0.781 |
| ML | LR(MACCS) | 0.439 / 0.774 | 0.334 / 0.798 | 0.413 / 0.837 | 0.197 / 0.771 | 0.240 / 0.643 | 0.191 / 0.723 | 0.180 / 0.752 |
| GNN | Ligandformer | 0.520 / **0.821** | 0.540 / 0.862 | 0.454 / 0.860 | 0.251 / 0.825 | 0.359 / 0.744 | 0.294 / 0.807 | 0.218 / 0.840 |
| GNN | GAT | 0.491 / 0.797 | 0.463 / 0.868 | 0.471 / 0.870 | **0.287** / 0.805 | 0.338 / 0.735 | 0.267 / 0.796 | 0.294 / **0.857** |
| GNN | GCN | 0.464 / 0.805 | 0.474 / 0.861 | 0.469 / 0.863 | 0.262 / 0.802 | 0.334 / 0.727 | 0.338 / 0.808 | 0.230 / 0.838 |
| GNN | GIN | 0.498 / 0.811 | 0.558 / **0.885** | 0.492 / 0.872 | 0.261 / 0.805 | 0.352 / 0.736 | 0.308 / 0.810 | 0.251 / 0.856 |
| Sequence | ChemBERTa | 0.538 / 0.799 | 0.558 / 0.857 | 0.480 / 0.855 | 0.223 / 0.806 | 0.331 / 0.724 | 0.362 / 0.806 | 0.238 / 0.764 |
| Sequence | ChemBERTa2 | **0.553** / 0.814 | 0.557 / 0.851 | **0.547** / **0.892** | 0.258 / **0.828** | 0.362 / 0.742 | 0.351 / 0.816 | **0.302** / 0.811 |
| Sequence | MoLFormer | 0.373 / 0.802 | 0.523 / 0.860 | 0.507 / 0.869 | 0.210 / 0.810 | 0.384 / **0.746** | 0.306 / 0.812 | 0.248 / 0.802 |
| LLM-SAR | GPT5.5-SAR | 0.053 / 0.509 | 0.041 / 0.450 | 0.287 / 0.714 | 0.181 / 0.752 | 0.198 / 0.640 | 0.075 / 0.657 | 0.077 / 0.680 |
| LLM-SAR | GPT5.5-SAR + knowledge | 0.113 / 0.707 | 0.117 / 0.675 | 0.333 / 0.762 | 0.204 / 0.762 | 0.217 / 0.624 | 0.080 / 0.658 | 0.083 / 0.709 |
| LLM-SAR | Opus4.7-SAR | 0.060 / 0.620 | 0.056 / 0.574 | 0.241 / 0.751 | 0.209 / 0.744 | 0.249 / 0.661 | 0.136 / 0.699 | 0.167 / 0.707 |
| LLM-SAR | Opus4.7-SAR + knowledge | 0.244 / 0.746 | 0.177 / 0.779 | 0.341 / 0.789 | 0.211 / 0.785 | 0.213 / 0.643 | 0.093 / 0.676 | 0.138 / 0.711 |

Table 11: Murcko scaffold 5-fold CV model-by-endpoint matrix for Tox21 stress response assays. Each endpoint entry reports PR-AUC / ROC-AUC. Rankings are computed independently for each metric within each endpoint: the first ranked value is bold and underlined, and the second and third ranked values are underlined.

| Family | Model | SR-ARE | SR-ATAD5 | SR-HSE | SR-MMP | SR-p53 |
|---|---|---|---|---|---|---|
| ML | ExtraTrees(ECFP4) | 0.375 / 0.725 | 0.212 / 0.792 | 0.243 / 0.735 | 0.568 / 0.816 | 0.299 / 0.792 |
| ML | ExtraTrees(ECFP6) | 0.376 / 0.726 | 0.209 / 0.790 | 0.243 / 0.728 | 0.562 / 0.814 | 0.290 / 0.776 |
| ML | ExtraTrees(RDKit desc.) | 0.445 / 0.783 | 0.243 / 0.835 | 0.317 / 0.758 | 0.711 / 0.901 | **0.337** / 0.832 |
| ML | ExtraTrees(MACCS) | 0.392 / 0.734 | 0.200 / 0.787 | 0.234 / 0.745 | 0.576 / 0.822 | 0.227 / 0.783 |
| ML | RF(ECFP4) | 0.396 / 0.728 | 0.219 / 0.807 | 0.259 / 0.740 | 0.576 / 0.818 | 0.306 / 0.787 |
| ML | RF(ECFP6) | 0.385 / 0.725 | 0.212 / 0.803 | 0.259 / 0.736 | 0.575 / 0.818 | 0.294 / 0.779 |
| ML | RF(RDKit desc.) | 0.450 / **0.785** | 0.221 / 0.818 | **0.318** / 0.766 | 0.711 / 0.900 | 0.336 / 0.836 |
| ML | RF(MACCS) | 0.412 / 0.741 | 0.220 / 0.812 | 0.262 / 0.757 | 0.598 / 0.832 | 0.247 / 0.810 |
| ML | GBDT(ECFP4) | 0.349 / 0.693 | 0.179 / 0.752 | 0.245 / 0.716 | 0.585 / 0.804 | 0.249 / 0.746 |
| ML | GBDT(ECFP6) | 0.332 / 0.676 | 0.164 / 0.741 | 0.217 / 0.680 | 0.557 / 0.793 | 0.228 / 0.723 |
| ML | GBDT(RDKit desc.) | 0.433 / 0.774 | **0.247** / 0.809 | 0.310 / 0.757 | **0.720** / **0.906** | 0.278 / 0.826 |
| ML | GBDT(MACCS) | 0.364 / 0.726 | 0.210 / 0.778 | 0.272 / 0.740 | 0.613 / 0.836 | 0.241 / 0.793 |
| ML | LR(ECFP4) | 0.298 / 0.650 | 0.147 / 0.691 | 0.131 / 0.620 | 0.441 / 0.712 | 0.193 / 0.687 |
| ML | LR(ECFP6) | 0.278 / 0.615 | 0.154 / 0.734 | 0.125 / 0.622 | 0.444 / 0.718 | 0.144 / 0.651 |
| ML | LR(RDKit desc.) | 0.393 / 0.745 | 0.157 / 0.776 | 0.225 / 0.751 | 0.578 / 0.839 | 0.225 / 0.795 |
| ML | LR(MACCS) | 0.319 / 0.706 | 0.151 / 0.762 | 0.233 / 0.725 | 0.548 / 0.822 | 0.189 / 0.783 |
| GNN | Ligandformer | **0.451** / 0.778 | 0.216 / 0.827 | 0.291 / 0.779 | 0.708 / 0.894 | 0.296 / 0.827 |
| GNN | GAT | 0.428 / 0.768 | 0.184 / 0.815 | 0.272 / 0.764 | 0.693 / 0.881 | 0.302 / 0.825 |
| GNN | GCN | 0.419 / 0.755 | 0.195 / 0.819 | 0.255 / 0.766 | 0.672 / 0.882 | 0.318 / 0.823 |
| GNN | GIN | 0.423 / 0.768 | 0.225 / **0.838** | 0.286 / **0.795** | 0.687 / 0.893 | 0.322 / **0.842** |
| Sequence | ChemBERTa | 0.392 / 0.747 | 0.222 / 0.777 | 0.249 / 0.762 | 0.646 / 0.866 | 0.237 / 0.778 |
| Sequence | ChemBERTa2 | 0.437 / 0.783 | 0.221 / 0.803 | 0.299 / 0.786 | 0.711 / 0.900 | 0.314 / 0.820 |
| Sequence | MoLFormer | 0.403 / 0.774 | 0.189 / 0.778 | 0.279 / 0.786 | 0.690 / 0.892 | 0.248 / 0.778 |
| LLM-SAR | GPT5.5-SAR | 0.271 / 0.617 | 0.061 / 0.615 | 0.097 / 0.570 | 0.307 / 0.566 | 0.105 / 0.632 |
| LLM-SAR | GPT5.5-SAR + knowledge | 0.335 / 0.682 | 0.063 / 0.624 | 0.116 / 0.608 | 0.419 / 0.709 | 0.124 / 0.677 |
| LLM-SAR | Opus4.7-SAR | 0.221 / 0.534 | 0.065 / 0.618 | 0.137 / 0.658 | 0.344 / 0.669 | 0.129 / 0.618 |
| LLM-SAR | Opus4.7-SAR + knowledge | 0.330 / 0.697 | 0.064 / 0.622 | 0.149 / 0.688 | 0.478 / 0.761 | 0.173 / 0.688 |

Table 12: Murcko scaffold 5-fold CV model-by-endpoint matrix for bioactivity-related classification endpoints. Each endpoint entry reports PR-AUC / ROC-AUC. Rankings are computed independently for each metric within each endpoint: the first ranked value is bold and underlined, and the second and third ranked values are underlined.

| Family | Model | EGFR | anti-TB H37Rv | antimalaria Pf. 3D7/Dd2 |
|---|---|---|---|---|
| ML | ExtraTrees(ECFP4) | 0.982 / 0.946 | 0.504 / 0.767 | 0.962 / 0.935 |
| ML | ExtraTrees(ECFP6) | **0.983** / **0.946** | 0.491 / 0.759 | 0.961 / **0.936** |
| ML | ExtraTrees(RDKit desc.) | 0.974 / 0.923 | 0.455 / 0.741 | 0.863 / 0.900 |
| ML | ExtraTrees(MACCS) | 0.970 / 0.916 | 0.410 / 0.717 | 0.940 / 0.916 |
| ML | RF(ECFP4) | 0.982 / 0.945 | 0.508 / **0.776** | **0.963** / 0.934 |
| ML | RF(ECFP6) | 0.982 / 0.944 | 0.495 / 0.762 | 0.960 / 0.928 |
| ML | RF(RDKit desc.) | 0.974 / 0.919 | 0.476 / 0.747 | 0.852 / 0.882 |
| ML | RF(MACCS) | 0.972 / 0.916 | 0.441 / 0.743 | 0.953 / 0.915 |
| ML | GBDT(ECFP4) | 0.971 / 0.913 | 0.460 / 0.734 | 0.949 / 0.921 |
| ML | GBDT(ECFP6) | 0.972 / 0.916 | 0.463 / 0.756 | 0.937 / 0.909 |
| ML | GBDT(RDKit desc.) | 0.968 / 0.902 | 0.461 / 0.761 | 0.883 / 0.914 |
| ML | GBDT(MACCS) | 0.961 / 0.885 | 0.402 / 0.716 | 0.925 / 0.901 |
| ML | LR(ECFP4) | 0.936 / 0.838 | 0.328 / 0.676 | 0.877 / 0.895 |
| ML | LR(ECFP6) | 0.945 / 0.858 | 0.338 / 0.704 | 0.892 / 0.890 |
| ML | LR(RDKit desc.) | 0.944 / 0.847 | 0.363 / 0.703 | 0.830 / 0.880 |
| ML | LR(MACCS) | 0.943 / 0.840 | 0.335 / 0.683 | 0.820 / 0.838 |
| GNN | Ligandformer | 0.975 / 0.921 | 0.470 / 0.772 | 0.903 / 0.925 |
| GNN | GAT | 0.973 / 0.915 | 0.500 / 0.763 | 0.911 / 0.923 |
| GNN | GCN | 0.974 / 0.918 | 0.486 / 0.768 | 0.905 / 0.918 |
| GNN | GIN | 0.976 / 0.925 | **0.511** / 0.770 | 0.916 / 0.925 |
| Sequence | ChemBERTa | 0.969 / 0.904 | 0.461 / 0.736 | 0.861 / 0.912 |
| Sequence | ChemBERTa2 | 0.974 / 0.918 | 0.498 / 0.759 | 0.921 / 0.924 |
| Sequence | MoLFormer | 0.979 / 0.932 | 0.504 / 0.763 | 0.914 / 0.929 |
| LLM-SAR | GPT5.5-SAR | 0.840 / 0.664 | 0.207 / 0.481 | 0.693 / 0.562 |
| LLM-SAR | GPT5.5-SAR + knowledge | 0.847 / 0.680 | 0.269 / 0.594 | 0.710 / 0.633 |
| LLM-SAR | Opus4.7-SAR | 0.842 / 0.657 | 0.230 / 0.476 | 0.884 / 0.779 |
| LLM-SAR | Opus4.7-SAR + knowledge | 0.909 / 0.766 | 0.325 / 0.636 | 0.904 / 0.847 |

### Supplementary Note 2b: Protocol Sensitivity of GNN and Sequence Model Readout

The primary optimal held-out readout reports the best held-out fold metric within fixed training settings for GNN and sequence models. To test whether the benchmark conclusions depend on this readout, we re-aggregated the same GNN and sequence model training logs using the fixed final-window readout, defined as the mean of the final three epochs. Classical ML and LLM-SAR results were unchanged.

Under the primary optimal held-out readout, classical ML accounts for 47.4% of strict family winners, sequence models for 28.8%, GNNs for 21.8% and LLM-SAR for 1.9%. Under the fixed final-window readout, the corresponding shares are 84.6%, 10.9%, 2.6% and 1.9%. This sensitivity analysis shows that strict winner shares depend on GNN and sequence model readout. It supports the manuscript's central conclusion that model scale alone does not order predictive performance and that classical ML is the most stable leading family, while GNN and sequence models remain endpoint dependent complements.

Table 13: Fold mean task and metric family winners by split and endpoint class under the fixed final-window readout. GNN and sequence models are read out as the mean of the final three training epochs; classical ML and LLM-SAR are unchanged from the primary comparison. Classification endpoints contribute PR-AUC and ROC-AUC winners; regression endpoints contribute MAE and Pearson winners. MAE uses the lowest value as the winner; all other metrics use the highest value.

| Split | Endpoint class | Task and metric entries | ML | GNN | Sequence | LLM-SAR | Total |
|---|---|---|---|---|---|---|---|
| Random | ADME related | 12 | 8 | 0 | 4 | 0 | 12 |
| | Toxicity related | 34 | 33 | 0 | 0 | 1 | 34 |
| | Bioactivity related | 6 | 6 | 0 | 0 | 0 | 6 |
| Murcko scaffold | ADME related | 12 | 8 | 2 | 2 | 0 | 12 |
| | Toxicity related | 34 | 30 | 0 | 3 | 1 | 34 |
| | Bioactivity related | 6 | 6 | 0 | 0 | 0 | 6 |
| Structure separated | ADME related | 12 | 6 | 2 | 4 | 0 | 12 |
| | Toxicity related | 34 | 30 | 0 | 3 | 1 | 34 |
| | Bioactivity related | 6 | 5 | 0 | 1 | 0 | 6 |
| Total | All endpoint classes | 156 | 132 | 4 | 17 | 3 | 156 |

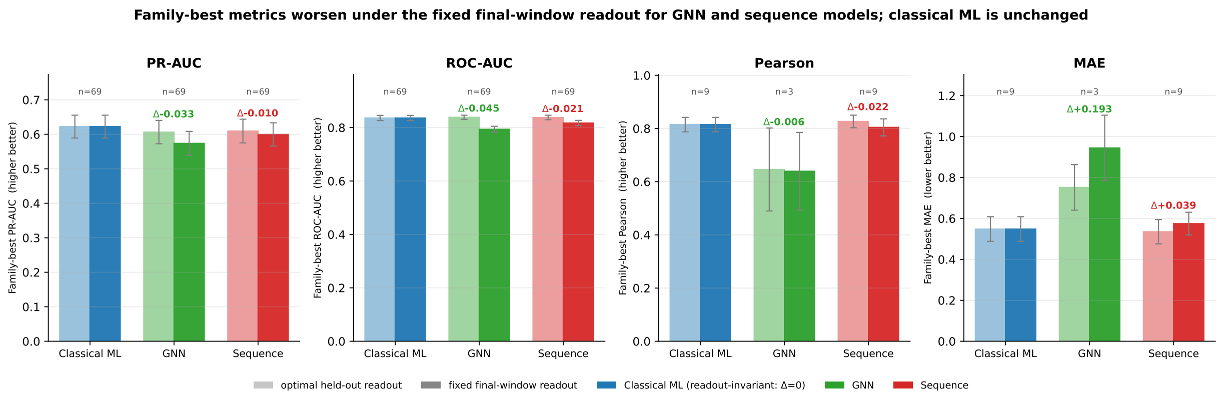


Figure 1: Sensitivity of GNN and sequence model family best metrics to readout definition, pooled across all endpoint and split entries. Each family shows two bars: the primary optimal held-out readout (light) and the fixed final-window readout (dark), with bars giving the mean family best metric and error bars the standard error across entries. Classical ML is unchanged by the readout definition, so its two bars are equal and serve as a readout-invariant baseline. The annotation over each neural family gives $\Delta = \text{fixed} - \text{optimal}$; for the higher-is-better metrics (PR-AUC, ROC-AUC, Pearson) a negative $\Delta$ indicates a lower value under the fixed final-window readout, whereas for MAE (lower-is-better) a positive $\Delta$ indicates a higher error. PR-AUC and ROC-AUC are computed from classification entries ($n = 69$ each), whereas MAE and Pearson are computed from the smaller set of regression entries ($n$ shown per panel).

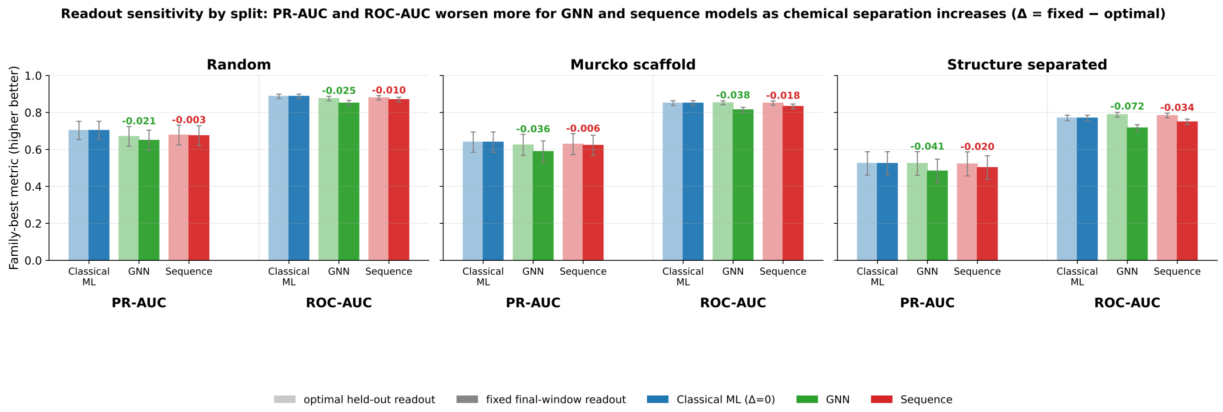


Figure 2: Readout sensitivity resolved by split protocol for the two classification metrics with full coverage (PR-AUC and ROC-AUC, $n = 23$ entries per metric and split group). Within each split panel, PR-AUC and ROC-AUC are grouped separately, and each family shows the optimal held-out readout (light) and fixed final-window readout (dark) family best mean with standard error; the annotation over each neural family is $\Delta = \text{fixed} - \text{optimal}$. Classical ML is readout-invariant. The fixed final-window decrease for GNN and sequence models grows with chemical separation (largest under structure separated CV), indicating that the strict first-place reassignments toward neural families under harder splits are sensitive to the held-out epoch readout. Regression metrics are omitted from this split-resolved view because GNN has no regression entries under random and Murcko scaffold CV.

### Supplementary Note 3: Paired Bootstrap Tests

Paired bootstrap tests use 1,000 resamples over the shared held out prediction pool. Classification resampling is stratified where possible, and Benjamini and Hochberg false discovery rate correction is applied within each level and metric. The main text reports two levels, family level and model level. Table 14 shows the coverage. The complete set of 1,404 paired bootstrap tests is provided in the supplementary data archive, with no skipped comparisons.

Table 14: Paired bootstrap test coverage by level. Family level compares best of family models on the same task across the four families. Model level compares the three pairwise contrasts among the top ranked, second ranked and third ranked models where available.

| Quantity | Family level | Model level |
|---|---|---|
| Pairs per entry | 6 | 3 |
| Classification entries | 69 | 69 |
| Classification tests per metric | 414 | 207 |
| Regression entries | 9 | 9 |
| Regression tests per metric | 54 | 27 |

### Supplementary Note 4: Top K Enrichment and Calibration

Top K enrichment and calibration are reported as supporting analyses for the corresponding main text subsections. The Top K analysis reports family best EF@1%, EF@5% and EF@10% by endpoint class; the calibration analysis reports family best Brier score and 10 bin ECE by endpoint class. Machine readable summaries are provided with the supplementary data. Table 15 summarises coverage. For antimalaria Pf. 3D7/Dd2 under Murcko scaffold CV, held out fold indices 1 and 4 contained only active compounds (49 and 33 molecules, respectively); EF@$K$ and calibration summaries for this split therefore use fold indices 0, 2 and 3.

Table 15: Top K enrichment and calibration coverage. Top K metrics include EF1%, EF5%, EF10% and top 50 / top 100 precision. Calibration metrics include Brier score and 10 bin expected calibration error (ECE). Rows are family best model records for the available split and task entries.

| Scope | Split and task entries | Top K rows | Calibration rows |
|---|---|---|---|
| Primary support set | 76 | 218 | 219 |

### Supplementary Note 5: Restricted Data Sensitivity Analysis

The primary infectious disease benchmark uses anti-TB H37Rv and antimalaria Pf. 3D7/Dd2 public activity datasets. Restricted institutional infectious disease panels are not included in the primary family winner denominator, the abstract percentages or the main endpoint matrices. They are retained only for restricted data sensitivity analysis, because their raw structural records require source provenance checks and release permission review before public redistribution.

Table 16: Restricted institutional infectious disease panels retained as supplementary sensitivity analyses. These panels are not used in the primary benchmark denominator.

| Endpoint | Activity definition | Records | Active | Inactive | Active rate |
|---|---|---|---|---|---|
| anti-TB H37Rv cellular activity | MIC $\leq 1\ \mu$M | 49,266 | 8,013 | 41,253 | 16.3% |
| antimalaria Pf. 3D7/Dd2 cellular activity | $EC_{50}$ $\leq$100 nM | 2,088 | 644 | 1,444 | 30.8% |

The restricted anti-TB H37Rv panel gives higher absolute discrimination values than the public anti-TB H37Rv core panel, as expected for a larger and partly institutional training distribution, but it does not change the qualitative interpretation of the main benchmark. Classical ML remains a strong family best baseline, GNN and sequence models remain close in several split and metric settings, and LLM-SAR remains below the trained model families in direct predictive performance.

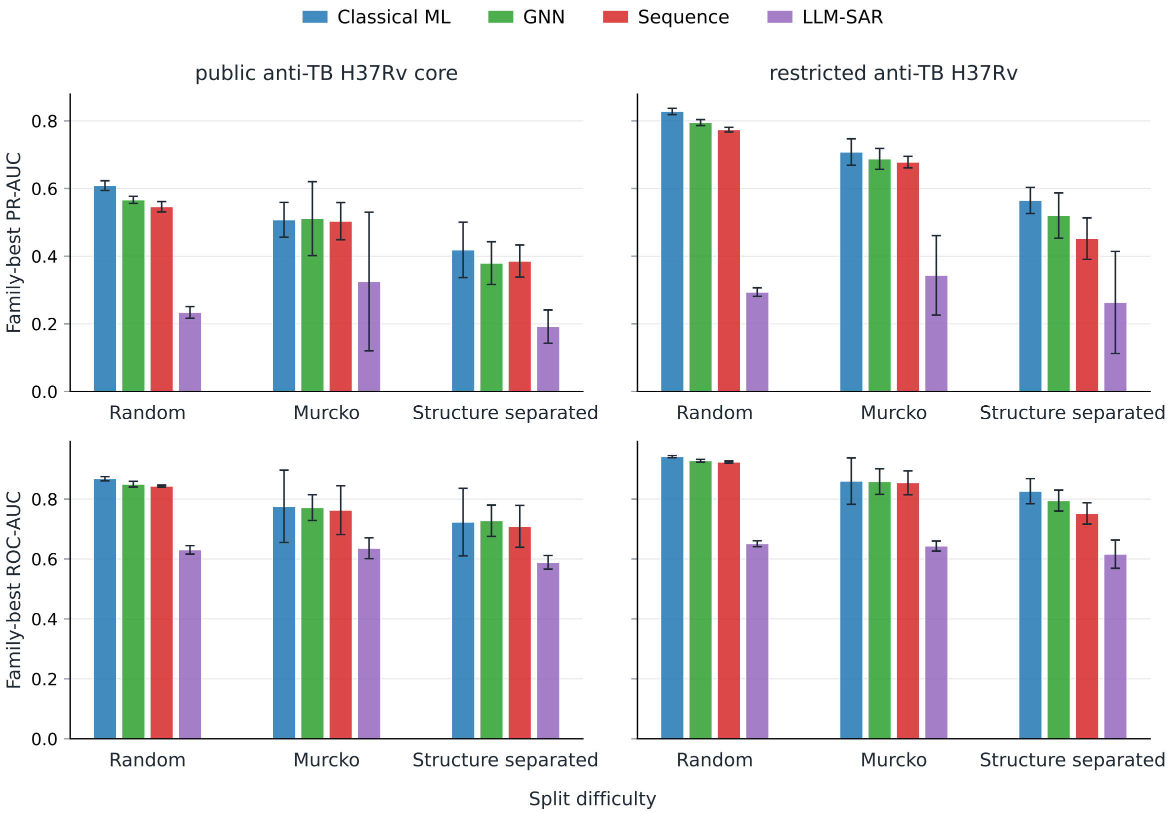


Figure 3: Restricted data sensitivity analysis for anti-TB H37Rv. Lines show family best fold mean PR-AUC and ROC-AUC across random, Murcko scaffold and structure separated 5-fold CV for the public anti-TB H37Rv core panel and the restricted institutional anti-TB H37Rv panel. Error bars show fold standard deviation.

Table 17: Restricted data sensitivity analysis for anti-TB H37Rv. Values are family best fold mean ± fold standard deviation for each split and metric. The restricted institutional panel is shown only as a supplementary sensitivity analysis and is not included in the primary family winner denominator.

| Dataset | Split | Metric | ML | GNN | Sequence | LLM-SAR | Leading family |
|---|---|---|---|---|---|---|---|
| restricted anti-TB H37Rv | Random | PR-AUC | RF(ECFP4) 0.828 ± 0.009 | GIN 0.795 ± 0.009 | ChemBERTa2 0.774 ± 0.007 | Opus4.7-SAR + knowledge 0.294 ± 0.013 | ML |
| restricted anti-TB H37Rv | Random | ROC-AUC | RF(ECFP4) 0.942 ± 0.004 | GIN 0.928 ± 0.005 | ChemBERTa2 0.924 ± 0.004 | Opus4.7-SAR + knowledge 0.651 ± 0.010 | ML |
| restricted anti-TB H37Rv | Murcko | PR-AUC | ExtraTrees(ECFP4) 0.708 ± 0.039 | GIN 0.688 ± 0.031 | ChemBERTa2 0.678 ± 0.017 | Opus4.7-SAR + knowledge 0.343 ± 0.118 | ML |
| restricted anti-TB H37Rv | Murcko | ROC-AUC | ExtraTrees(ECFP4) 0.860 ± 0.077 | GIN 0.858 ± 0.043 | ChemBERTa2 0.854 ± 0.040 | Opus4.7-SAR + knowledge 0.643 ± 0.017 | ML |
| restricted anti-TB H37Rv | Structure separated | PR-AUC | RF(ECFP4) 0.565 ± 0.039 | GAT 0.520 ± 0.067 | ChemBERTa2 0.452 ± 0.061 | GPT5.5-SAR + knowledge 0.263 ± 0.151 | ML |
| restricted anti-TB H37Rv | Structure separated | ROC-AUC | RF(ECFP4) 0.826 ± 0.042 | GAT 0.795 ± 0.035 | ChemBERTa 0.752 ± 0.036 | GPT5.5-SAR + knowledge 0.616 ± 0.047 | ML |
| public anti-TB H37Rv core | Random | PR-AUC | RF(ECFP4) 0.609 ± 0.014 | GIN 0.567 ± 0.010 | ChemBERTa2 0.546 ± 0.015 | Opus4.7-SAR + knowledge 0.234 ± 0.017 | ML |
| public anti-TB H37Rv core | Random | ROC-AUC | RF(ECFP4) 0.868 ± 0.007 | GIN 0.850 ± 0.009 | ChemBERTa2 0.843 ± 0.003 | Opus4.7-SAR + knowledge 0.631 ± 0.014 | ML |
| public anti-TB H37Rv core | Murcko | PR-AUC | RF(ECFP4) 0.508 ± 0.051 | GIN 0.511 ± 0.109 | MoLFormer 0.504 ± 0.055 | Opus4.7-SAR + knowledge 0.325 ± 0.205 | GNN |
| public anti-TB H37Rv core | Murcko | ROC-AUC | RF(ECFP4) 0.776 ± 0.121 | Ligandformer 0.772 ± 0.043 | MoLFormer 0.763 ± 0.081 | Opus4.7-SAR + knowledge 0.636 ± 0.035 | ML |
| public anti-TB H37Rv core | Structure separated | PR-AUC | ExtraTrees(ECFP4) 0.419 ± 0.082 | Ligandformer 0.380 ± 0.063 | MoLFormer 0.386 ± 0.047 | Opus4.7-SAR + knowledge 0.192 ± 0.049 | ML |
| public anti-TB H37Rv core | Structure separated | ROC-AUC | RF(ECFP6) 0.723 ± 0.113 | GAT 0.728 ± 0.052 | ChemBERTa2 0.709 ± 0.070 | Opus4.7-SAR + knowledge 0.589 ± 0.023 | GNN |

The restricted infectious disease LLM-SAR sensitivity analysis also supports the main interpretation. SAR knowledge derived from training folds improves anti-TB H37Rv ranking metrics and improves Opus4.7-SAR on the restricted antimalaria Pf. 3D7/Dd2 panel, but these rule based variants remain lower than the best trained model families.

Table 18: Restricted data sensitivity analysis for LLM-SAR with and without training fold SAR knowledge. Values are fold mean metrics for the restricted institutional infectious disease panels.

| Endpoint | Metric | GPT5.5-SAR | GPT5.5-SAR + knowledge | Opus4.7-SAR | Opus4.7-SAR + knowledge |
|---|---|---|---|---|---|
| anti-TB H37Rv cellular activity | PR-AUC | 0.203 | 0.263 | 0.205 | 0.261 |
| anti-TB H37Rv cellular activity | ROC-AUC | 0.548 | 0.616 | 0.514 | 0.609 |
| antimalaria Pf. 3D7/Dd2 cellular activity | PR-AUC | 0.374 | 0.361 | 0.336 | 0.358 |
| antimalaria Pf. 3D7/Dd2 cellular activity | ROC-AUC | 0.522 | 0.522 | 0.478 | 0.521 |

Only aggregate derived results are provided for these restricted panels. The corresponding summary files are provided in the reproducibility package rather than the underlying restricted structures:

- `reproducibility_code_data/data/restricted_data_manifest/restricted_infectious_disease_dataset_summary.csv`
- `reproducibility_code_data/data/restricted_data_manifest/restricted_infectious_disease_llm_sar_metrics.csv`
- `reproducibility_code_data/data/figure_source_data/supplementary_figure3_restricted_data_sensitivity.csv`

These files allow readers to inspect whether the restricted institutional sensitivity analyses support the same qualitative trends without requiring access to the underlying restricted structures.
